# Metagenomic Sequencing Detects Respiratory Pathogens in Hematopoietic Cellular Transplant Patients

**DOI:** 10.1101/102798

**Authors:** C Langelier, MS Zinter, K Kalantar, GA Yanik, S Christenson, B Odonovan, C White, M Wilson, A Sapru, CC Dvorak, S Miller, CY Chiu, JL DeRisi

## Abstract

**RATIONALE:** Current microbiologic diagnostics often fail to identify the etiology of lower respiratory tract infections (LRTI) in hematopoietic cellular transplant recipients (HCT), which precludes the implementation of targeted therapies.

**OBJECTIVES:** To address the need for improved LRTI diagnostics, we evaluated the utility of metagenomic next generation sequencing (mNGS) of bronchoalveolar lavage (BAL) to detect microbial pathogens in HCT patients with acute respiratory illnesses.

**METHODS:** We enrolled 22 post-HCT adults ages 19-69 years with acute respiratory illnesses who underwent BAL at the University of Michigan between January 2012 and May 2013. mNGS was performed on BAL fluid to detect microbes and simultaneously assess the host transcriptional response. Results were compared against conventional microbiologic assays.

**MEASUREMENTS & MAIN RESULTS:** mNGS demonstrated 100% sensitivity for detecting respiratory microbes (human metapneumovirus, respiratory syncytial virus, *Stenotrophomonas maltophilia*, human herpesvirus 6 and cytomegalovirus) when compared to standard testing. Previously unrecognized LRTI pathogens were identified in six patients for whom standard testing was negative (human coronavirus 229E, human rhinovirus A, *Corynebacterium propinquum* and *Streptococcus mitis*); findings were confirmed by independent PCR and 16S rRNA sequencing. Relative to patients without infection, patients with infection had increased expression of immunity related genes (p=0.022) and significantly lower diversity of their respiratory microbiome (p=0.017).

**CONCLUSIONS:** Compared to conventional diagnostics, mNGS enhanced detection of pathogens in BAL fluid from HCT patients. Furthermore, our results suggest that combining unbiased microbial pathogen detection with assessment of host gene biomarkers of immune response may hold promise for enhancing the diagnosis of post-HCT respiratory infections.

## INTRODUCTION

Lower respiratory tract infections (LRTI) are a leading reason for hospitalization and mortality in hematopoietic cellular transplant (HCT) recipients (1, 2). This problem is underscored by autopsy studies showing that previously undetected pulmonary pathogens may contribute to death in 30% of HCT recipients (3, 4). Despite this, the etiologic pathogens remain unidentified in most cases of LRTI, due largely to the limitations of current microbiologic tests in terms of sensitivity, speed and breadth of assay targets (5).

Diagnosis of LRTI in HCT recipients is particularly challenging due to high rates of non-infectious inflammatory conditions such as graft versus host disease (GVHD) that can drive pulmonary inflammation, induce fever and mimic infection (6). Furthermore, the diagnostic yield of traditional tests is reduced in HCT patients due to antimicrobial prophylaxis, reduced antibody titers, and infections from uncommon opportunistic microorganisms (7).

The limitations of current microbiologic tests drive excessive use of empiric broad-spectrum antimicrobials, which potentiates the emergence of drug resistance and increases risk of *Clostridium difficile* infection (8). In some situations, empiric regimens may lack activity against an underlying microbe, leading to treatment failure, disease progression, and consequent adverse outcomes (5, 9). Furthermore, in transplant patients, concern for GVHD may compel clinicians to initiate empiric immunosuppressive agents that could inadvertently exacerbate disease in the presence of an unrecognized infection (10).

Previously, microarray approaches proved useful for broadening the scope of pathogens detectable in a single LRTI assay; some microarrays have even incorporated conserved sequences from all known viruses, allowing detection of novel viral strains (11). Metagenomic next-generation sequencing (mNGS) inherently offers enhanced diagnostic capabilities by providing a culture-independent, comprehensive measurement of the microbial composition of clinical samples (12, 13). This technology permits the simultaneous detection of bacterial, viral and fungal pathogens without introducing bias associated with fixed-target PCR or serologic assays (11, 12, 14). By capturing both microbial and human RNA, mNGS also permits simultaneous transcriptional profiling of the host immunologic response, which can provide complementary insight regarding the presence and type of infection (15, 16). Due to the clear need for enhanced respiratory pathogen diagnostics in HCT recipients, we undertook this study examining the utility of host/pathogen mNGS for detection of LRTI pathogens in HCT patients hospitalized with acute respiratory illnesses.

## MATERIALS AND METHODS

### Subjects

This retrospective study evaluated sequentially enrolled adult HCT recipients who underwent bronchoscopy with bronchoalveolar lavage (BAL) at the University of Michigan Medical Center between January 25^th^, 2012 and May 20^th^, 2013. All patients met established clinical definitions of community-acquired or hospital-acquired pneumonia and had two or more of the following: cough, fever, chills, dyspnea, pleuritic chest pain, crackles, or bronchial breathing; in addition, an opacity, infiltrate or nodules on chest radiograph was required (17-19). In some cases, patients had other equally probable non-infectious explanations for respiratory symptoms. All subjects had routine microbiological testing of BAL fluid as part of standard of care diagnostic workup for respiratory illness. Patients consented to permit banking of surplus BAL specimens in a biorepository at -80°C in accordance with University of Michigan IRB protocol HUM00043287. De-identified samples were then transported to UCSF for mNGS.

### Clinical Microbiologic Testing

During the period of study enrollment, standard of care clinical microbiologic diagnostic testing for respiratory pathogens included bacterial, mycobacterial and fungal cultures, CMV culture, *Aspergillus* galactomannan assay, multiplex PCR for influenza A/B, respiratory syncytial virus (RSV) and human metapneumovirus (HMPV), human herpesvirus-6 (HHV-6) PCR, and silver stain for *Pneumocystis jiroveci*. Additional studies included nasophayngeal (NP) swab PCR for influenza A/B, RSV, and HMPV, blood cultures, serum PCR for herpes simplex virus (HSV) types 1 and 2, varicella-zoster virus (VZV), Epstein-Barr virus (EBV), cytomegalovirus (CMV) and HHV-6, *Clostridium difficile* toxin stool PCR, and select additional studies on BAL per the clinical judgment of the treating physicians.

### Metagenomic Next-Generation Sequencing

Total nucleic acid was extracted from 250µl of patient BAL fluid using bead-based lysis and the Zymo Viral DNA/RNA Kit (Zymo Research). RNA and DNA were isolated separately using DNAse or RNAse, respectively, and the former was reverse transcribed to generate complementary DNA (cDNA). DNA and cDNA then underwent adapter addition and barcoding using the Nextera system (Illumina). Depletion of abundant sequences by hybridization (DASH) was employed to selectively deplete human mitochondrial cDNA, thus enriching for microbial transcripts (20). The final RNAseq and DNA sequencing (DNAseq) libraries underwent 135 nucleotide paired-end Illumina sequencing. Methodology is described further in the online data supplement.

### Pathogen Detection Bioinformatics

Detection of microbial pathogens leveraged a custom bioinformatics pipeline to discriminate pathogens from background microbial sequences in clinical samples. This methodology is detailed in the Online Data Supplement (21). Briefly, this analytical approach involved first filtering for quality and complexity, next extracting sequences aligning to the human and Pan troglodytes genomes (NCBI GRC h38 and UCSC PanTro4) using STAR (22) and then subsequently removing reads aligning to non-fungal eukaryotes and phage using Bowtie2 (23). The identities of the remaining reads were determined by querying the NCBI nucleotide (nt) and non-redundant protein (nr) databases using GSNAP-L.

To mitigate spurious taxonomic assignments due to laboratory background contaminants, we calculated a standard Z-score for each microbial genus relative to a control group of 17 “no-template” water-only controls (detailed in Table E1) derived from sequencing runs performed in our laboratory over the past 2 years (24). Z-scores were calculated based on unique reads aligned to either NCBI nt or nr, per million sequence input reads (rpM_nt_, rpM_nr_).

The respiratory tract is not a sterile environment, and as such, we devised a ranking score to aid in the stratification of the inherently complex alignment data. This metric involved multiplying the number of reads aligned per million reads sequenced by the sum of the nt and nr Z-scores for each microbial genus identified in each patient: score = rpM_nt_ × (Z_nt_ + Z_nr_). To further improve the specificity of microbial assignments, we required 1) microbes to be detected in both nucleotide (nt) and protein (nr) database alignments; 2) microbes to have a Z_nt_ and Z_nr_ > zero and 3) bacteria and fungi detected by RNAseq to also have detectable genomic sequence by DNAseq. In addition, we permitted DNA viruses to be detected on only DNAseq and RNA viruses to be detected on only RNAseq. For each patient, any microbes that passed these requirements were then processed using the following decision tree, summarized in Figure 1.

**Figure 1.**
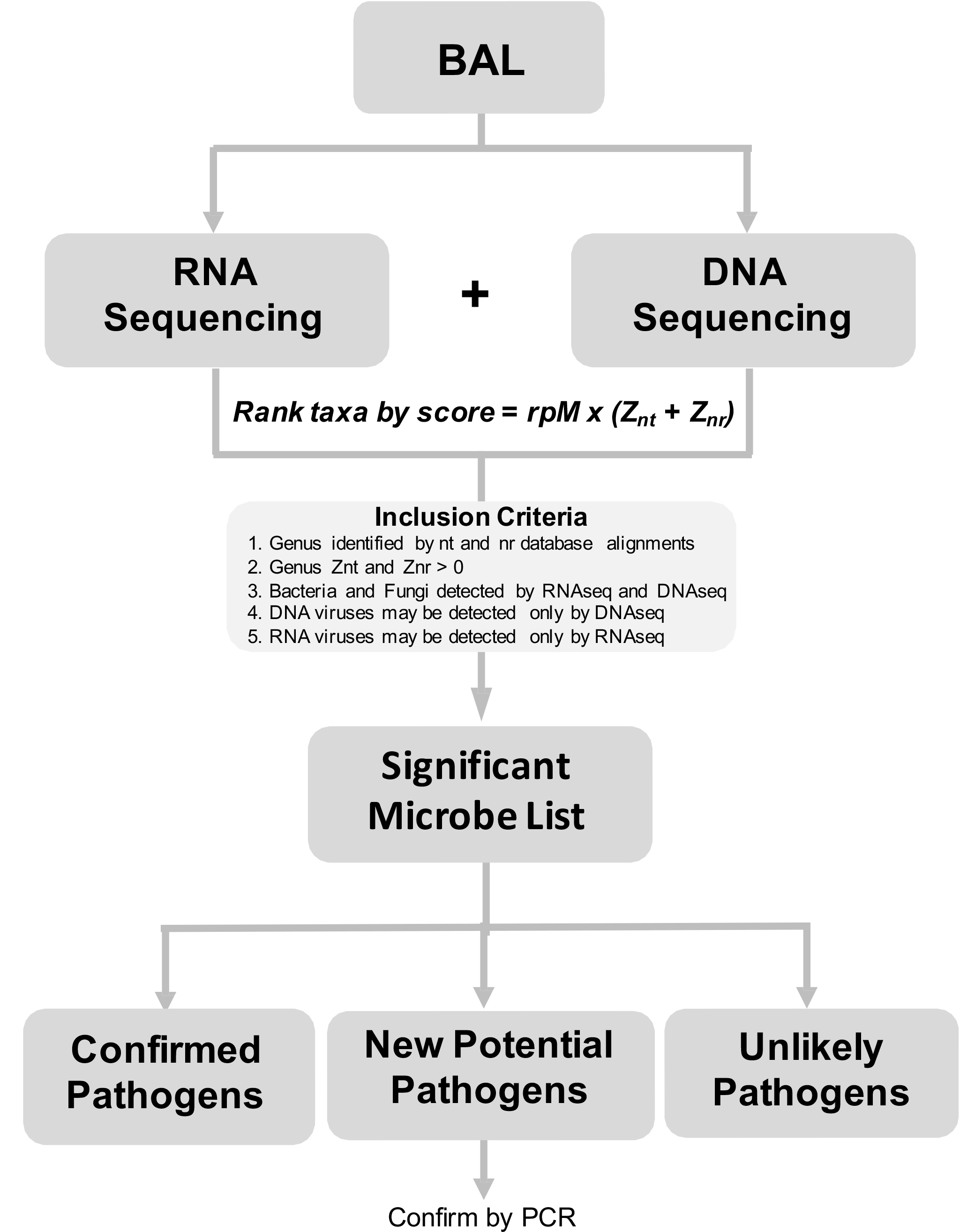
Overview of Pathogen Detection Workflow. Total nucleic acid extracted from BAL fluid of HCT recipients underwent DNA and RNA sequencing. A custom bioinformatics pipeline simultaneously identified pathogens and assayed the human transcriptome. Pathogens meeting inclusion and exclusion criteria were ranked by microbial significance score [Score = rpM_nt_ × (Z_nt_ + Z_nr_)]. To further improve the specificity of microbial assignments: we required 1) microbes to be detected in both nt and nr database alignments, 2) microbes to have a Z_nt_ and Z_nr_ greater than zero, and 3) bacteria and fungi detected by RNAseq to also have detectable genomic sequence by DNAseq. For each patient, any microbes that passed these requirements were then classified as either 1) confirmed pathogens, 2) new potential pathogens or 3) unlikely or uncertain pathogens.

### Definitions

Microbes were considered **confirmed pathogens** if a) both clinical testing and mNGS identified the microbe, b) there existed literature evidence of microbial pathogenicity in the lungs, and c) the microbe score was as least two-fold greater than that of any other microbe of the same type (virus, bacteria or fungus) identified in the patient (17). Microbes were considered **new potential pathogens** if a) mNGS alone identified the microbe, b) there existed literature evidence of microbial pathogenicity in the lungs, and c) the microbe score was two-fold greater than any other microbe of the same type (virus, bacteria or fungus) in the patient. New potential pathogens were confirmed by independent specific PCR testing for viruses and 16s rRNA gene Sanger sequencing for bacteria. Finally, microbes were considered **unlikely or uncertain pathogens** if they a) lacked robust literature evidence of microbial pathogenicity in the lungs, b) had a score that was less than two-fold that of the top-ranking microbe, suggesting a polymicrobial sample, c) determined to be clinically insignificant by the treatment team, or d) were a DNA virus of uncertain pathogenicity present in low abundance (< 5rpM).

### Microbial diversity

Alpha diversity of the respiratory microbiome in each subject was assessed using the Simpson Diversity Index (SDI) and compared between patients with a) confirmed pathogens, b) unlikely or uncertain pathogens and c) new potential pathogens using the nonparametric Wilcoxon Rank Sum test (25).

### Host gene expression analyses

RNA transcripts aligning to the human genome were captured by our computational pipeline as described above. Cumulative sum scaling normalization of protein-coding gene transcripts was carried out (26) and genes expressed in fewer than 30% of samples, or as outliers in only 10% of samples, were removed. Using a supervised approach, pathways related to immune functionality were selected *a priori* from the Molecular Signatures Database (27) and compared in terms of total normalized expression between subjects using the nonparametric Wilcoxon Rank Sum test as described in the online data supplement. These pathways contained gene biomarkers associated with antiviral response, interferons α, β, and γ, IL-6/JAK/STAT5 signaling and adaptive immunity, and have all been strongly associated with the immune response to infection (28, 29).

## RESULTS

### Cohort characteristics and clinical outcomes

We enrolled 22 HCT recipients hospitalized for acute respiratory symptoms aged 19-69 years at a median 356 days post-transplant, as summarized in Table 1. Their most common transplant indications were leukemia (n=12) and lymphoma (n=8). The majority of patients received allogeneic HCT (n=20) and/or myeloablative conditioning (n=20), as summarized in Table E2. The median absolute neutrophil count (ANC) at the time of BAL was 3.8/µL (IQR 2.1-5.4) and engraftment was achieved at the time of BAL in all patients. Acute GVHD was present in three subjects and chronic GVHD was present in 13 subjects. At the time of BAL, 15 subjects were receiving immune modulating agents.

**Table 1:** Clinical and Microbiologic Data. Abbreviations: <u>Radiography</u>: B/L, bilateral; LUL, left upper lobe; RUL, right upper lobe. Chest CT was obtained for all subjects except 1, 3, 4, 8, 9 and 14 who received chest X-rays. <u>Antimicrobials</u>: ACV, acyclovir; CIDV cidofovir; CPM, cefepime; FOSC, foscarnet, FLUC, fluconazole; GCV, ganciclovir; MICA, micafungin; POSA, posaconazole; RIBV, ribavirin; TMP-SMZ, trimethoprim-sulfamethoxazole; TOBR, tobramycin; vACV, valacyclovir; VANC, vancomycin; VORI, voriconazole. <u>Respiratory Support</u>: HFNC, high flow nasal cannula; IPPV; invasive positive pressure ventilation; NC, nasal cannula. *Clinicians concluded that CMV in subject 37 was not the principal cause of respiratory disease.

| Patient ID | Symptoms and Duration | Chest Radiography | Standard BAL Microbiology | Top Scoring Microbes by mNGS | Non-BAL Microbiology | Antimicrobial Prophylaxis | Antimicrobial Treatment | GVHD | Respiratory Support | Cause of death |
| --- | --- | --- | --- | --- | --- | --- | --- | --- | --- | --- |
| **Pathogens identified by conventional and NGS diagnostics** |  |  |  |  |  |  |  |  |  |  |
| 1 | Fever, hypoxia<br>x 3 days | RLL, LLL infiltrate | *HMPV* | *HMPV* | *HMPV* (NP), *C. difficile* (stool) | ACV, LEVO, MICA, TMP-SMZ | CFPM, VANC | aGVHD, cGVHD | NC | Relapse, day 253 |
| 5 | Cough, dyspnea, fever<br>x 3 days | Multifocal nodules, bronchiectasis | *Stenotrophomonas maltophilia* | *S. maltophilia* + *HRV-A* | Negative | ACV, LEVO, TMP-SMZ, VORI | TMP-SMZ, VANC | aGVHD, cGVHD | NC | GVHD, day 462 |
| 8 | Dyspnea<br>x 12 days | B/L GGO | *RSV* | *RSV* | CMV (blood) | ACV, LEVO, PENT, POSA | CIDV, FOSC, GCV, VANC, (IVIG) | No | HFNC | Relapse, day 265 |
| 9 | Cough<br>x 3 days | RUL, RML, LUL GGO, nodules | Negative | *HMPV* | *HMPV* (NP) | ACV, TMP-SMZ | None | cGVHD | None | alive |
| 10 | Cough, fever<br>x 4 days | Bibasilar airspace disease | *RSV* | *RSV* | *RSV* (NP) | FLUC, LEVO, vACV | CFPM, RIBV, (IVIG) | aGVHD, cGVHD | NC | GVHD, day 3024 |
| 36 | Dyspnea, fever<br>x 1 day | Diffuse septal thickening, GGO | *HHV-6* | *HHV-6* | HHV-6 (blood) | ACV, FLUC, LEVO, VORI | FOSC | No | None | alive |
| **New potential pathogens identified by NGS** |  |  |  |  |  |  |  |  |  |  |
| 6 | Cough, dyspnea<br>x 3 days | B/L nodules | Negative | *HCOV-229E* | Negative | TMP-SMZ, vACV, VORI | CFPM, TOBR, VANC | cGVHD | None | alive |
| 7 | Dyspnea<br>x 1 day | B/L perihilar opacities | Negative | *HCOV-229E* | Negative | ACV, FLUC, LEVO, TMP-SMZ | CFPM, VANC, (IVIG) | aGVHD, cGVHD | IPPV x 4 days | GVHD, day 96 |
| 13 | Cough, dyspnea<br>x 2 days | LLL opacities, air trapping | Negative | *Corynebacterium propinquum* | Negative | ACV, FLUC, LEVO, TMP-SMZ | CFPM, VANC | No | IPPV x 3 days | Idiopathic respiratory failure, day 51 |
| 14 | Cough, fever<br>x 2 days | LLL opacities | Negative | *HRV-A* | Negative | TMP-SMZ | None | cGVHD | None | alive |
| 18 | Cough, fever<br>x 2 days | RML, RLL atelectasis | Negative | *HRV-A* | *S. epidermidis* (blood) | FLUC, LEVO, vACV | CFPM, VANC | n/a | None | alive |
| 19 | Dyspnea<br>x 4 wks | RUL, LUL reticular opacities | Negative | *Streptococcus mitis* | Negative | ACV, LEVO, VORI | VANC | aGVHD, cGVHD | None | alive |
| **Microbes of uncertain or unlikely pathogenicity** |  |  |  |  |  |  |  |  |  |  |
| 3 | Fever<br>x 2 days | B/L infiltrates | Negative | *Flavobacterium sp.* | Gram positive rod (blood) | ACV, LEVO, TMP-SMZ, VORI | VANC | No | None | GVHD, day 267 |
| 11 | Cough, fever<br>x 2 days | LLL opacities | Negative | Negative | Negative | ACV, FLUC, LEVO, TMP-SMZ | CFPM | aGVHD, cGVHD | None | alive |
| 22 | Dyspnea, rigors<br>x 1 day | RUL, LUL peribronchial thickening, GG | Negative | *Torque teno virus* | Negative | ACV, LEVO, TMP-SMZ, VORI | CFPM | cGVHD | NC | alive |
| 23 | Cough, dyspnea, fever<br>x 3 wks | B/L GG, lymphadenopathy | Negative | Negative | EBV viremia | ACV, LEVO, MICA, TMP-SMZ, VORI | CFPM | aGVHD | IPPV x 10 days (until death) | No Autopsy, day 10 |
| 24 | Dyspnea, fever<br>x 1 wk | B/L perivascular and airspace GG | Negative | *Pyrenophora sp.* | Negative | ACV, TMP-SMZ, VORI | CFPM, TOBR | aGVHD, cGVHD | NC | alive |
| 25 | Cough, dyspnea, fever<br>x 4 wks | Diffuse bronchiectasis, multifocal GG nodules | Negative | *WU Polyoma Virus* | Negative | LEVO, VORI, TMP-SMZ | VANC | aGVHD, cGVHD | NC | alive |
| 31 | Dyspnea<br>x 2 wks | B/L GG, bronchiectasis | Negative | *Prevotella* | Negative | ACV, FLUC, LEVO, TMP-SMZ | CFPM | cGVHD | NC | alive |
| 34 | Cough, dyspnea<br>x 2 wks | Multifocal nodules, opacities, air trapping | Negative | *Torque teno virus* | Negative | ACV, TMP-SMZ | None | cGVHD | None | alive |
| 35 | Cough, dyspnea<br>x 2 days | Multifocal GG, tree-in-bud opacities | Negative | Negative | Negative | ACV, FLUC, LEVO | VANC | aGVHD, cGVHD | NC | alive |
| 37 | Cough, dyspnea<br>x 8 days | B/L GG | CMV* | CMV* | Negative | TMP-SMZ | None | aGVHD, cGVHD | None | Alive |

Fever was present in 13 of 22 (59%) of patients, all patients met systemic inflammatory response syndrome criteria, and no patients had septic shock. Twelve patients required supplemental oxygen, three required mechanical ventilation and all survived their illnesses except for one subject who died 10 days after BAL due to lymphoma relapse. Seven of the 22 (32%) subjects died between 51 and 3024 days post-BAL and their primary causes of death were GVHD (n=3), relapsed malignancy (n=2), idiopathic respiratory failure (n=1), and unknown (n=1).

### Antimicrobial use

All study subjects received antimicrobial prophylaxis prior to symptom onset and bronchoscopy. Empiric broad spectrum antibiotics were used in 16 of the 22 (73%) patients including vancomycin (n=4), cefepime (n=4), vancomycin + cefepime (n=5), cefepime + tobramycin (n=1), and vancomycin + cefepime + tobramycin (n=1) (Table 1). Pathogen-targeted antimicrobials were used in three subjects (14%). Four patients (15%) received no antimicrobials aside from pre-existing prophylactic agents.

### Clinical Microbiologic Findings

Standard of care clinical BAL diagnostics performed at the study hospital (described above in Methods) identified microbes in seven of 22 (32%) patients, of which six were thought to represent etiologic pathogens by the treating physicians. These included HMPV (n=2), RSV (n=2), HHV-6 (n=1) and *Stenotrophomonas maltophilia* (n=1). CMV was identified by shell vial culture in subject 37 but thought to represent incidental carriage because of symptom resolution in the absence of intervention prior to the return of testing.

### Next Generation Sequencing Findings

An average of 49 million paired-end sequencing reads were generated from each BAL sample, of which <1% were microbial. Sequencing statistics and pipeline output for each patient are described in Table E3. mNGS identified all seven microbes found by standard clinical diagnostics, demonstrating 100% sensitivity for pathogen detection. In total, RNAseq identified 10 RNA viruses, all of which have established pathogenicity in the lungs, and the RNA intermediates of five DNA viruses that have uncertain pulmonary pathogenicity. DNAseq identified the genomes of these same five DNA viruses as well as five others including CMV and HHV-6, which have been associated with pneumonitis. mNGS captured entire viral genomes for five patients at an average read depth of 3500-fold, as shown in Figure E1. Bacteria known to exist contextually as either pathogens or commensals, including *Stenotrophomonas maltophilia, Streptococcus mitis*, and *Corynebacterium propinquum* were identified, as were genera not typically considered pathogenic (30-32). Specificity of mNGS could not be assessed because this cohort lacked patients with proven non-infectious respiratory illnesses.

### Relationship Between Respiratory Microbial Diversity and Detection of Pathogens

Loss of diversity within respiratory tract microbial communities has been proposed as an ecological marker of infection (33, 34). We thus evaluated alpha diversity of actively replicating microbes identified by RNAseq using the SDI and found that subjects with confirmed pathogens had significantly lower diversity relative to patients with only microbes of unlikely or uncertain pathogenicity (0.34, IQR 0.15-0.64, n=6 vs. 0.92, IQR 0.86-0.93, n=10, p=0.017, Figure 2 and Table E4). Reduced diversity was also observed if subjects with confirmed or potential new pathogens were compared together against those without (0.41, IQR 0.20-0.55, n=12 vs. 0.92, IQR 0.86-0.93, n=10, p<0.001).

**Figure 2.**
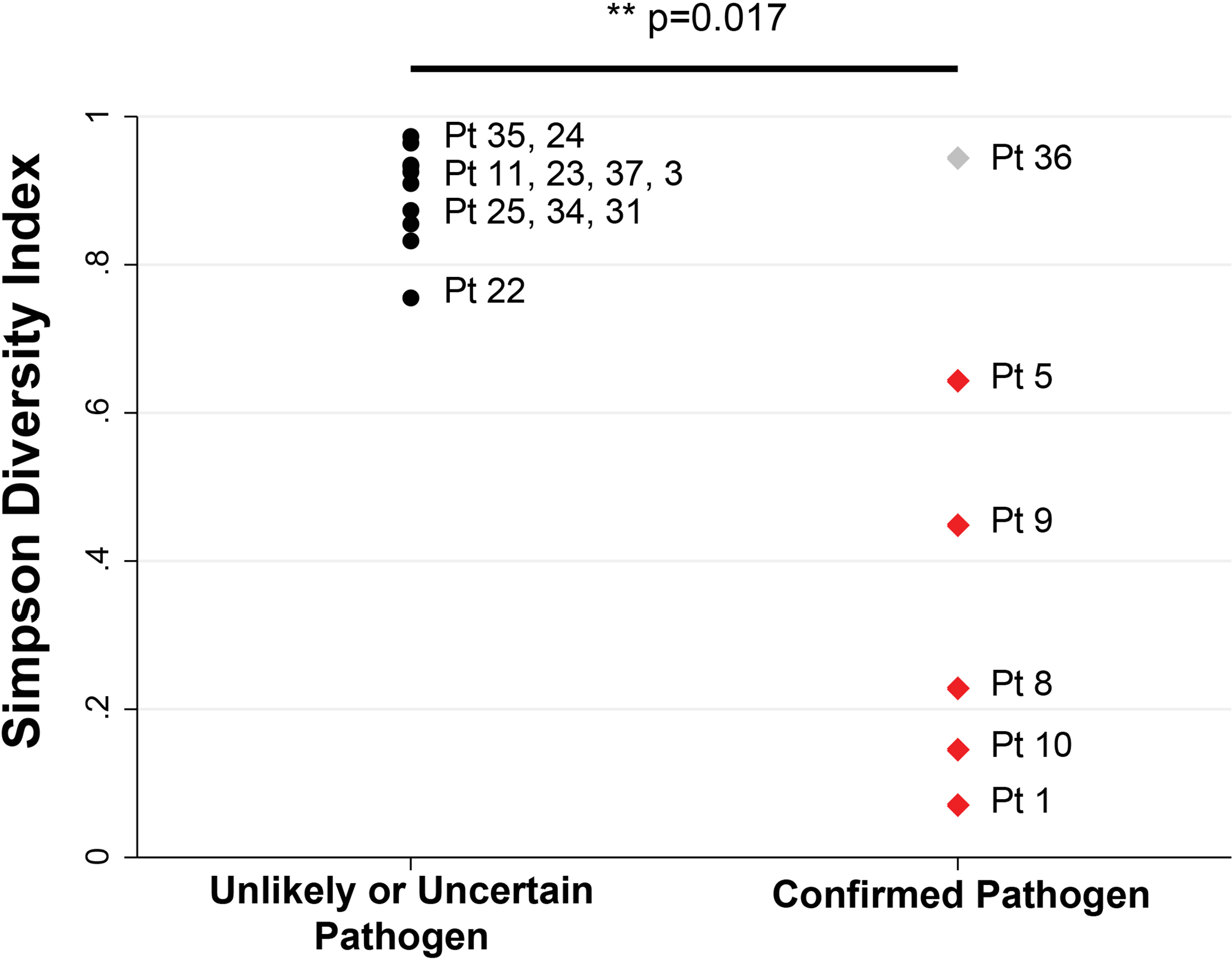
BAL Microbial Diversity is Inversely Associated with Presence of a Transcriptionally Active Respiratory Pathogen. Each data point represents a single patient for whom the Simpson Diversity Index (SDI) is plotted on the y-axis. Subjects are grouped according to confirmed pathogen (red triangles) vs. unlikely or uncertain pathogen (black circles). Patients with confirmed pathogens had significantly lower diversity relative to patients with only microbes of unlikely pathogenicity (0.34, IQR 0.15-0.64, n=6, vs. 0.92, IQR 0.86-0.93, n=10, p=0.017). Raw data are listed in **Table E4**.

### Analysis of Host Gene Expression

We hypothesized that gene expression from respiratory fluids obtained at the site of infection would distinguish patients with LRTI from those with non-infectious respiratory diseases. To test this idea, we evaluated *a priori* selected gene biomarkers of innate and adaptive immune responses using the total normalized expression of all genes. Given suspected between-subject immunologic response heterogeneity due to differences in infection type and degree of immunologic compromise, we created a composite metric based on the sum normalized gene expression of all biomarkers. We found significantly increased expression in patients with confirmed LRTI pathogens versus those without (94.9, IQR 93.8-105.6, n=6 vs. 33.1, IQR 20.7-75.1, n=7, p=0.022), as shown in Figure 3 and Table E5. When patients with confirmed or potential new pathogens were compared together against all others, the relationship trended towards significance (94.6, IQR 76.5-105.6, n=11 vs. 33.1, IQR 20.7-75.1, n=7, p=0.0634), and reached significance (p = 0.014) if a subject with a relatively lower abundance of HRV-A transcripts was instead grouped with patients found to have microbes of unlikely or uncertain pathogenicity. Raw gene counts are provided in the additional online supplemental data table.

**Figure 3.**
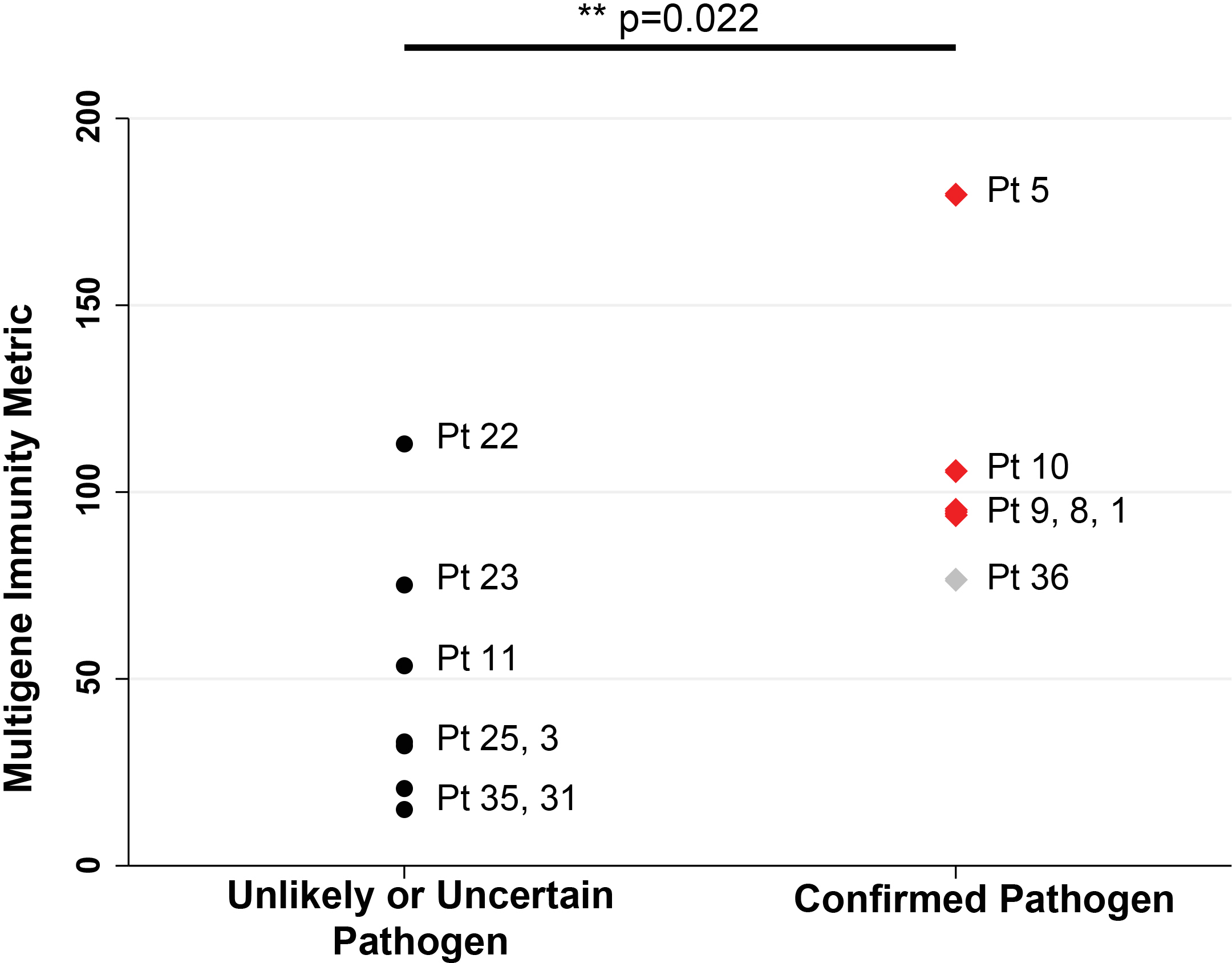
Expression of a Host Immune Response Multi-Gene Metric Correlates with Detection of LRTI Pathogens. Each data point represents a single patient for whom the composite immune response gene metric is plotted on the y-axis. Subjects are grouped according to confirmed pathogen (red triangles) vs. or unlikely or uncertain pathogen (black circles). Patients with confirmed pathogens had significantly higher multi-gene metric relative to patients with only microbes of unlikely pathogenicity (33.1, IQR 20.7-75.1, n=7 vs. 94.9, IQR 93.8-105.6, n=6, p=0.022). Raw data are listed in Table E5.

## DISCUSSION

In this proof of concept study, we demonstrate that mNGS can simultaneously detect pathogens and the host response in HCT patients with acute respiratory illnesses. When benchmarked against the conventional standard of care testing performed during the time of study enrollment, mNGS demonstrated 100% sensitivity for microbial detection and permitted identification of potential new viral and bacterial pathogens in six of 15 (40%) subjects with otherwise negative testing. We observed that the presence of respiratory pathogens was characterized by increased expression of host immune response genes and lower airway microbial diversity. Our results suggest that simultaneous mNGS detection of respiratory pathogens, the host’s immunologic response and the airway microbiome may provide complementary data that could inform the clinical significance of a detected pathogen.

Amongst the cases with negative clinical testing, mNGS identified six patients harboring transcriptionally active microbes with recognized respiratory pathogenicity (35-40). With respect to viruses, HCOV-229E was identified in two patients and HRV-A was identified in three patients, one of whom was also culture-positive for *Stenotrophomonas maltophilia*. Notably, HRV and HCOV were not represented on the multiplex PCR panel used at the study hospital; similar limited panels are still in use at many hospitals today. Unlike rapid antigen or multiplex PCR assays, mNGS is not limited to a fixed number of pre-specified targets on a panel, obviating the need to order multiple independent diagnostic tests. Furthermore, mNGS can detect entire viral genomes, as illustrated in Figure E1, an attribute that can enable genotyping, detection of resistance mutations, and epidemiologic tracking of disease outbreaks (41).

With respect to previously unidentified bacterial pathogens, mNGS detected *Streptococcus mitis*, an oropharyngeal microbe known to cause bacteremia and acute respiratory distress in HCT recipients, as a potential pulmonary pathogen in one patient (40, 42). *Corynebacterium propriniquum*, one of the few virulent *Corynebacterium* species associated with LRTI, comprised the majority of microbial transcripts in another subject’s BAL fluid (43). Both patients received empiric vancomycin, which has activity against *Streptococcus and Corynebacteria spp*., and recovered from their acute respiratory illnesses. Findings were independently confirmed at the genus level by 16s rRNA gene sequencing.

Ten patients with negative conventional testing were found to have microbes of unlikely or uncertain pathogenicity. Notably, each of these patients had potential alternative explanations for their respiratory symptoms. While one of these patients had bacteremia/sepsis, the remaining nine had acute and/or chronic GVHD, underlining the importance of non-infectious alloreactive inflammation in post-HCT pulmonary complications.

mNGS identified several viruses with DNA genomes, however only five of these also had well-defined evidence of active replication marked by detectable RNA transcripts (HSV, human papilloma virus and torque teno viruses (TTV, n=3). TTV was the most commonly detected DNA virus, a finding consistent with prior reports demonstrating an increased prevalence of this presumptively innocuous constituent of the human virome in immunosuppressed patients (44). Herpesviridae genomic DNA in the absence of viral transcripts was identified in five patients and included HHV-6 (n=1), CMV (n=2), HSV (n=1) and EBV (n=1). One subject, who died 10 days post-BAL due to relapsed lymphoma, had EBV DNA detected in the setting of EBV viremia. WU polyoma virus was detected by DNA sequencing another case, and while this virus has been associated with respiratory infections in immunocompromised patients, evidence supporting its role as a pathogen is lacking (45).

CMV was identified by both viral culture and mNGS in one subject, however this finding was considered unrelated to the patient’s respiratory illness by the treating physicians. Notably, while CMV was detected by DNAseq, no RNA transcripts were identified, suggesting that it may have represented incidental carriage as opposed to a transcriptionally active pathogen (46). This patient was also found to harbor *Pseudomonas fluorescens*, a described colonizer of the human respiratory tract (47), as well as several other taxa representing common constituents of the oropharyngeal or respiratory microbiome, including *Rothia, Prevotella*, and *Actinomyces spp*., as detailed in Table E3.

### Lung microbiome diversity and the host immune response as biomarkers of infection

Because asymptomatic carriage of respiratory pathogens is well described (48, 49), establishing biomarkers of genuine infection is critical for determining whether a given microbiologic finding is clinically significant. Our findings suggest that respiratory tract microbial diversity may be such a biomarker. Specifically, we found that patients with LRTI pathogens had significantly lower alpha diversity versus those without (p=0.017, Figure 2, Table E5), presumably reflecting dominance of actively replicating pathogens (33, 34).

Our results also demonstrate that both mNGS and conventional methods identify clinically significant and insignificant microbes, and emphasize the need to assess the impact of a given microbiologic finding in the context of a patient’s immunologic response (5). Expression of a multi-gene immune response metric was significantly increased in patients with confirmed respiratory pathogens relative to those without, suggesting that despite significant and heterogeneous states of immune suppression, HCT recipients still exhibited immunologic biomarkers of active infection (p=0.022, Figure 3).

Amongst subjects with potential new pathogens, we observed that two of the three HRV-A positive subjects demonstrated the lowest expression of this immune response metric, while the remaining subject, who was co-infected with HRV-A and *Stenotrophomonas maltophilia*, had one of the highest values. This is consistent with prior reports demonstrating that HRV can induce a broad range of clinical disease severity, and that viral-bacterial co-infection can increase the severity of disease (48, 49). Amongst subjects with confirmed pathogens, the patient with clinically suspected HHV-6 pneumonitis was notably an outlier in terms of reduced immune metric expression and elevated airway diversity. The respiratory pathogenicity of HHV-6 is controversial, and our paired mNGS pathogen data demonstrating absence of viral transcripts but detectable genomic DNA support a less virulent role for HHV-6 in this patient (50).

Broadspectrum antibiotic overuse is frequently driven by suspicion for occult pathogens missed by standard diagnostic testing. For instance, patient 1, whose conventional testing identified only HMPV, received empiric vancomycin and cefepime due to suspected occult bacterial infection and subsequently developed *C. difficile* colitis. mNGS confirmed the presence of HMPV and also demonstrated absence of bacterial pathogens in this patient. The theoretical availability of our mNGS findings during the actual period of patient hospitalization potentially could have informed more targeted antimicrobial use in the 18 study subjects who received broad spectrum antibiotics in the absence of detectable bacterial pathogens by mNGS.

While these results are highly encouraging, this proof-of-concept study suggests many routes for further improvement. First, this study was limited by a relatively small sample size, only one case of culture-positive bacterial LRTI, no fungal infections, and no subjects with clinically diagnosed non-infectious airway diseases. Future studies with larger cohorts will be needed to validate the sensitivity and specificity of mNGS for LRTI diagnosis in this population. Second, our limited sequencing depth did not yield the human transcriptome coverage that would be desired for robust differential gene expression analyses, although we were able to rigorously evaluate a composite metric of immunity genes. Future larger studies are needed to identify and validate gene classifiers that can distinguish LRTI from non-infectious airway diseases in HCT recipients.

## CONCLUSIONS

Here we leverage continued improvements in metagenomic sequencing to expand the capabilities of LRTI diagnostics in HCT recipients with acute respiratory illnesses. We demonstrate that compared to current microbial diagnostics, mNGS has a greater capacity for detecting microbes and an ability to couple pathogen detection with simultaneous profiling of the host response and the airway microbiome. We suggest several ways in which future studies may improve on this proof-of-concept study, including validation in a larger prospective cohort of HCT recipients and other high-risk populations.

